# Danish profile of soft rot *Pectobacteriaceae*; A three-year field sampling study proving several clonal clades of soft rot isolates across diverse locations implicating a common origin

**DOI:** 10.64898/2026.05.11.724364

**Authors:** Julie Stenberg Pedersen, Laura Milena Forero Junco, Anna Struebel, Magnus M. Rothgardt, Frank Hille, Charles M.A.P. Franz, Birgit Jensen, Tue Kjærgaard Nielsen, Witold Kot, Chayan Roy, Alexander Byth Carstens, Lars Hestbjerg Hansen

## Abstract

Soft rot *Pectobacteriaceae* (SRP) are among the most economically important plant pathogenic bacteria and are especially known to be problematic in potato production. The epidemiology of disease transmission has been investigated for almost a century, and several aspects have been highlighted as plausible infection routes. However, it is generally accepted that the major source of disease is the latently infected mother tuber, but several parameters are still influencing disease prevalence including contaminated equipment, soil water status as well as temperature. Management of the disease is limited to hygiene practices, dry storage and seed certification systems but several studies have also proven biocontrol agents such as bacteriophages (phages) as promising tools. Despite the severity of SRP on potato production, little is known about the genetic diversity of SRPs in Denmark, and since only few isolates are available, the possibility to design a broadly effective phage cocktail is limited. Here we describe a three-year field study utilizing an agri-citizen science approach where Danish farmers provided symptomatic potato plants or tubers, together with metadata such as date, location, potato variety and origin. By using whole genome sequencing (Illumina and Nanopore) together with metadata we were able to investigate and monitor the epidemiological disease spread across the country using 103 complete genomes, sampled across all three years. In this study we provide epidemiological evidence of disease origins and a suite of phages that could be used as a biocontrol tool for early disease intervention. Our results revealed several clonal clades across diverse locations (SNPs < 20) which strongly indicate common origin. A total of 17 Pectobacterium phages were tested and did target > 80% of clonal clades. Based on the clonality across the soft rot isolates we propose the possibility to set in early on using phages targeting strains relevant for soft rot development, with the possibility of a surveillance program together with customizing the phage preference.

## INTRODUCTION

Soft rot *Pectobacteriaceae* (SRP) (formerly known as soft rot *Erwinia*) includes the genera *Pectobacterium* and *Dickeya*, both widely known as some of the most important bacterial plant pathogens within agriculture (Mansfield et al., 2012). SRPs are able to cause disease in 50% of the angiosperm plant order, but are especially economically impactful in potato production, both for seed and ware production, where it has been estimated that the overall yearly loss in Europe is up to 42 M euro (Dupuis et al., 2021; B. Ma et al., 2007). The epidemiology of SRP in potato production has been debated for almost a century, where several key components for understanding the underlying transmission routes have been proposed (Czajkowski et al., 2011). Already in 1926 observations in Minnesota pointed towards a transmission route driven by a symbiotic relationship between soft rot bacteria and the fly *Delia platura*. More recent work by Rossman et al. showed *Delia* species recovered from potato fields to carry SRPs but also laboratory-reared species to carry SRPs, indicating SRPs as a natural part of their microbiome, which may indicate *Delia* species to be a possible source of infection (Leach, 1926; Rossmann et al., 2018). Since 1926 numerous studies have investigated possible transmission routes for SRPs and latently infected mother tubers were later proposed as the disease source, and the seed certification system (based on multiplication on disease free stem cuts), was initiated in Scotland in 1967 (The Scottish Classification Scheme). In a six-year survey published in 1980, Pérombelon et al. provided some indication of disease transmission via mechanical equipment, based on observation from soft rot development in disease free stem cuts propagated across multiple farms (Pérombelon et al., 1980). Later, Pérombelon described how latently infected mother tubers may result in diseased plants as well as correlating with higher disease incidences in later generations, affected by both soil water status and temperature (Pérombelon, 1992). Other studies have suggested both aerosols during rain and irrigation water as plausible transmission routes, and SRPs have been isolated from river waters (Ben Moussa et al., 2022; Elphinstone & Pérombelon, 1987; Pédron et al., 2020). Jonkheer et al. did nonetheless indicate *Pectobacterium* isolates from river waters to be avirulent when tested (Jonkheer et al., 2021). It is now widely accepted that the major disease source is the latently infected mother tuber, however possible transmission routes which still are recognized as critical regarding disease development are both insects, aerosols, irrigation water, soil and equipment (van der Wolf et al., 2021).

Disease management is currently based on seed certification systems, high hygiene and optimized storage conditions, as no high throughput methods using physical treatments are possible and chemical treatments with antibiotics are banned due to the environmental risks (van der Wolf et al., 2021). A wide range of biocontrol agents have been proposed as alternative treatment mechanisms, such as bacteriophages (phages) (Charkowski, 2018). Although it was a century ago that Koons and Kotila first demonstrated phages capable of protecting against tissue soft rot in tubers and carrots, recent research continues to underscore their potential as effective biocontrol agents in potato productions (Beňo et al., 2022; A. B. Carstens et al., 2019; Coons & Kotila, 1925; Petrzik et al., 2023; Zaczek-Moczydłowska et al., 2020). In Denmark the Danish meristem program (seed certification based on a meristem of the tuber) was initiated in 1977 and completely implemented for all professional seed potato growers in 1986. However, in later years there has been reported an increase of soft rot incidences in the early generations of seed potatoes (Nielsen, 2013; personal communication with Danish plant consultants).

This study utilized an agri-citizen science approach by collaborating with Danish plant consultants and farmers to disseminate sampling kits and engaging Danish potato farmers across the country. Over a period of three years Danish potato farmers were invited to send potato tubers and plants symptomatic with soft rot or blackleg, which were used for isolation of SRPs. All isolates were used for whole-genome sequencing to investigate the epidemiological disease spread across the country. Furthermore, a bacteriophage isolation campaign where some isolates were used for isolation of bacteriophages was initiated. These were tested against the collection of isolated SRPs in relation to their biocontrol potential.

## METHODS AND MATERIALS

### Sample and metadata collection year 2021 – 2023

In collaboration with plant consultants across Denmark sample collection kits were handed out to Danish potato farmers during spring (February to May) in 2021, 2022 and 2023 (Suppl. fig. 1A). Sample kits for potato farmers included a project - and sample technique description for the potato farmer, plastic bags for the potato plant or tuber displaying blackleg or soft rot symptoms (Suppl. fig. 1B), together with an accompanying document for the potato farmer to fill out and include in the sample sending (Suppl. fig. 1A). Together with the potato plant or tuber displaying disease systems the farmer would fill out the accompanying document with area (county), district (municipality) and potato variety. When receiving the sample, the date of the sample was noted together with the rest of the metadata.

### Bacterial isolation

#### Sample isolation year 2021 – 2023

Bacteria were isolated as shortly described in previous work (Pedersen et al., 2024). The isolation method was adapted from Zaczek-Moczydłowska et al., 2019, using crystal violet pectin single layer (CVP) described by Hélias et al., 2012 with Dipecta (Agdia-Bioford) as the pectin source. Potato plants or tubers displaying soft rot symptoms send by Danish farmers (received between May and November in the years 2021, 2022 and 2023) were used for isolation. Samples were taken with a sterile loop inside the macerated tissue (in stem or tuber) and diluted in 1.6 ml sterile Mili-Q water. Samples were vortexed for 30 sec. and incubated for at least 5 min. at room temperature (RT, ∼ 25 °C) and vortexed again for 30 sec. 100 µl of the plant extract were spread on CVP plates using a Drigalski spatula, where the same spatula was used for another four successive CVP plates to dilute the sample. CVP plates were incubated for 48 h at 28 °C. Cavity forming colonies were re-streaked onto new CVP plates and incubated at 48 h at 28 °C. Single cavity forming colonies were then streaked onto Lysogeny Broth (LB) plates with 1.5 % agar and incubated between 24 – 48 h at 28 °C. Overnight (ON) cultures from single colonies were done using LB and incubated for 24 h at 28 °C. 15 % glycerol stocks of ON cultures were prepared and stored at - 80 °C for further use.

#### Focused sample isolation year 2023

To investigate whether seed potatoes would be latently carrying the same bacterial strains as appearing in plants or tubers displaying soft rot symptoms in the following season, three lots of seed potatoes were tested in 2023. The three lots were received from a Danish potato company. The lot numbers were as follows: 3009, 2296 and 2264. No tubers showed soft rot symptoms at the time of handling. 10 tubers from each lot were randomly chosen, and incubated in a sterile beaker with 500 ml sterilized tap water ON at 150 rpm at RT for 24 h. The tubers were taken from the water and examined in a laminar flow hood. Tubers displaying soft rot like symptoms (macerated tissue) the day after were sampled from and tested on CVP plates like described above and purified isolates named as J lot number_sample (e.g. J2264_1. To enrich any *Pectobacterium* species in the sample we also used the enrichment media media LEM_AG366,_ prepared as described by Hélias et al., 2012. The remaining water (incubated with tubers ON) was centrifuged at 10,000 g for 5 min and the supernatant was poured off except for 100 ml. The remaining 100 ml was used to resuspend the pellet for at least 90 min at 150 rpm at RT. 100 ml of the sample was then added to 900 ml of the enrichment media LEM_AG366_. The same procedure as described in the section above was used with 100 µl of the ON enrichment as input. Purified isolates were named as J lotnumberLEM.

### DNA extraction

For all isolates 1 ml of ON culture was used as input for the Genomic Mini AX Bacteria (A&A Biotechnology) following the manufacturers protocol and the DNA was dissolved in sterile water (Sigma Aldrich). Afterwards the Clean-Up Concentrator kit (A&A Biotechnology) was used for DNA cleanup following the manufacturers protocol. The DNA concentration was measured using a Qubit 2 instrument and the Qubit^TM^ Broad Range Assay Kits (ThermoFisher) and diluted in sterile water (Sigma Aldrich) according to the chosen library building protocol.

### Library building and genome sequencing

#### Bacterial isolates year 2021

All 2021 isolates were sequenced using both Illumina and Nanopore platforms, as described in previous work (Pedersen et al., 2024). Sequencing libraries for Illumina paired-end sequencing were prepared using the Nextera XT DNA library Kit (Illumina), and indexed DNA was sequenced on Illumina NextSeq500 platform using the Mid Output kit v2.5 (300 cycles). Sequencing libraries for Nanopore sequencing were prepared using the Rapid Barcoding Kit v14 (Oxford Nanopore Technologies), and indexed DNA was sequenced using the Nanopore MinION. To complete all genome assemblies some genomes were re-sequenced using the same barcoding kit and the Nanopore PromethION.

#### Bacterial isolates year 2022 – 2023

All 2022 – 2023 isolates were sequenced using Nanopore. Sequencing libraries were prepared using the Rapid Barcoding Kit v14 (Oxford Nanopore Technologies), and indexed DNA was sequenced using the Nanopore PromethION.

#### Bacterial isolates used for single nucleotide polymorphism analysis

For some bacterial isolates from 2022 and 2023 used for single nucleotide polymorphism (SNP) analysis, DNA was also sequenced using Illumina. Library preparations were done using the Hackflex protocol (Gaio et al., 2022) with the Illumina DNA prep kit (Illumina). Briefly, 10 µl Bead-Linked Transposomes (BLT) (1:50) and 25 µl of Tagmentation buffer 1 (TB1) were added to 10 µl of purified DNA, mixed and incubated for 15 min at 55 °C and held for 10 °C. 10 µl of Tagment Stop Buffer (TSB) was then added to the sample and incubated for 15 min at 37 °C and held at 10 °C. Samples were then transferred to a magnetic rack until clear suspension (∼3 min). The supernatant was removed and discarded. The samples were moved to a cold block and the PCR Master Mix (10 µl of 5x PCRBIO HiFi Buffer (PCR-BIOSYSTEMS), 0.5 µl PCRBIO HiFi Polymerase (PCRBIOSYSTEMS) and 29.5 µl sterile water (Sigma Aldrich)) was added together with the addition of 5 µl of each adaptor i5 and i7 and mixed. The following conditions were used for the PCR cycle: initial denaturation at 95 °C for 1 min, following 12 cycles of denaturation (95 °C for 15 sec), annealing (62 °C for 30 sec), extension (68 °C for 2 min) and afterwards final extension (68 °C for 1 min) and held at 10 °C. Samples were moved to the magnetic rack until clear suspension (∼5 min) and 50 µl of the supernatant was transferred to a new tube. 50 µl of pure water (Sigma Aldrich) and 50 µl of Cleanup Ampure XP Beads were added to the sample and incubated at RT (∼25 °C) for 5 min then placed on the magnetic rack until clear suspension (∼5 min). 150 µl of the supernatant was transferred to new tubes and 16.5 µl of AMPure XP beads added to the sample and incubated at RT for 5 min. Samples were then placed on the magnetic rack until clear solution (∼5 min) and the supernatant was discarded. 180 µl of 80% ethanol was added and incubated for 30 sec before supernatant was removed, the wash step was repeated. The beads were airdried for less than 5 min on the magnetic stand and the samples were removed from the magnetic stand and 22 µl of nuclease free water was added. Samples were incubated at RT for 2 min and placed on the magnetic stand until clear solution (∼2 min) and 20 µl of the supernatant transferred to a new tube. Indexed DNA was sent off and sequenced at Novogene on Illumina NovaSeq X Plus Series (PE150).

### Genome assembly and annotation

Genome assemblies for 2021 isolates were done using raw reads from Illumina and Nanopore sequencing runs with Unicycler (v0.5.0) (Wick et al., 2017). Bakta (v1.9.3) was used to annotate all genomes from 2021 isolates (Schwengers et al., 2021). Genome assemblies for 2022 and 2023 isolates were done using raw Oxford Nanopore reads from barcoded samples, which were processed using the MOSAIC Snakemake workflow from the GitHub repository lauramilena3/MOSAIC, main branch commit c8694b2. The workflow was executed with Snakemake v7.18.2. Read quality was assessed with nanoQC v0.9.4 and NanoStat v1.5.0. Adapters were removed with Porechop v0.2.4 using --discard_middle, and reads were filtered with NanoFilt v2.8.0 using -q 10, -l 1000, --maxlength 10000000, --headcrop 50, and --tailcrop 50. Filtered reads were assembled with Flye v2.9.5 using Nanopore high-quality/metagenomic settings. Assemblies were polished with Medaka v1.7.2 using the r941_min_high_g360 model, followed by two rounds of Racon v1.4.20 using Minimap2 v2.23 alignments. Assembly statistics were generated with QUAST v5.3.0. Assemblies were annotated using Bakta v1.11.0 with Bakta database v6.0. Genome completeness and contamination were estimated with CheckM-genome v1.2.5 using the checkm_data_2015_01_16 database. To ensure all assemblies were complete we did only include assemblies which met the following criteria for downstream analysis: genomic contigs <12 and n50 values >5kb, resulting in 103 complete genomes (of 144 isolates). To reorient all complete assemblies and share same start site we used Dnaapler (v0.8.0) (Bouras et al., 2024). Taxonomic classification was performed using GTDB-Tk v2.3.2 with GTDB release R214.

### Phylogenetic analysis and genomic characterization

The phylogenetic tree for the 103 complete genomes was created using PhyloPhlAn (v3.2.1) with options ‘- diversity high’ and ‘-fast’ (Asnicar et al., 2020). SNP calling was done for clonal clades in CLC Genomics Workbench (v22) (QIAGEN) using map reads to reference, default parameters (earliest isolate was chosen as reference in each clade except for *P. brasiliense* clade I, where it was the second isolation), following basic variant detection with all parameters set to default except for the following; ‘min cov. 10’, ‘min. count 2’, ‘min freq. 70%’ and ‘read direction filter 20%’. Visualizations of whole genome alignments (using contig 1 if >1 contig was present in the assembly), within all clonal clades were done using Dot_plot_like_in_blast (v1.6) (Schelkunov, 2024).

### Phage classification

#### Phage isolation, DNA extraction and sequencing

Phage isolation and purification was done as described in previous work for all phages except for Pectobacterium phage mrpotatohead and mspotatohead (Pedersen et al., 2024). Briefly, the organic waste sample was centrifuged for 10,000 × g for 10 min at 4 °C and filtered through 0.45 µm PVDF Syringae filters (Fisherbrand™) before use. All hosts for phage isolation are noted in supplementary table 1 together with all phage isolates. All hosts were grown in LB broth ON at 200 rpm at 28 °C. Phages were isolated and purified using the standard double agar overlay as described elsewhere (Pedersen et al., 2024). Pectobacterium phage mrpotatohead and mspotatohead were isolated from a washed tuber, described in (Pedersen, Carstens, Bollen, et al., 2025). Prior to isolation the tuber was soaked in sterile tap water for 30 min and dried in a laminar flow bench placed in a petridish for 15 min and afterwards twisted and placed in another petridish for 5 min, following incubation for 24h at 5 °C with air ventilation in a plastic box with premade wholes (radius of ∼ 0.3 cm). After 24h the tuber was cut into four with a sterile knife and incubated in a tuber with 20 ml of SM buffer (100 mM NaCl, 8 mM MgSO_4_·7H_2_O, 50 mM Tris-Cl pH 7.5) for 1h at 200 rpm, with rotation every 15 min. The SM solution was then centrifuged at 10,000 × g for 10 min at 4 °C and filtered through 0.22 µm PVDF Syringae filters (Fisherbrand™) and used for the standard double agar overlay as above, with host *Pectobacterium brasiliense* (J36_21). DNA extraction was done as described elsewhere for phages (A. Carstens et al., 2018). NEBNext® Ultra™ II FS DNA Library Prep Kit for Illumina (New England Biolabs) was used for library preparation for all phages, except for library preparation of phage mrpotatohead and mspotatohead which were prepared using Nextera XT DNA Library Preparation Kit (Illumina). Indexed DNA were sequenced using the Nextseq 500 platform with the Mid Output kit v2.5.

#### Phage assembly, annotation and comparative genomics

metaSPADES (v3.14.1) (Nurk et al., 2017) was used to assemble phage genomes from raw reads for all phages, except for phage mrpotatohead and mspotatohead which were assembled using CLC Genomics Workbench v22. Pharokka (v1.7.0) (Bouras et al., 2023) was used for genome annotation and VIRIDIC (Moraru et al., 2020) for calculating and visualizing the intergenomic similarity score for all versus all including previously isolated phages (Pappous (acc. no. PQ008973.1), Taid (acc. no. PQ008975.1), Abuela (acc. no. OP748251.2), Ymer (acc. no. OP748252.2), Koroua (acc. no. PQ008972.1), Amona (acc. no. Q008971.1), Sabo (acc. no. PQ008974.1), mrpotatohead, mspotatohead and Mimer (acc. no. ON872163.2)) (Pedersen, Carstens, Bollen, et al., 2025; Pedersen et al., 2024, 2026), phage Nepra (acc. no. NC_048704.1) (A. B. Carstens et al., 2019).

#### Phage transmission electron microscopy

Transmission Electron Microscopy (TEM) was done for a subset of phages as described in previous work (Pedersen et al., 2024). Briefly, phage lysates were pipetted on 100-mesh copper grids coated with carbon. Adsorption was allowed for 20 min and the grids were washed twice with deionized water and negatively stained with 2% uranyl acetate. Electron micrographs were generated on a Talos L120C transmission electron microscope (ThermoFisher Scientific, Eindhoven, The Netherlands) using a 4 k x 4 k Ceta camera (ThermoFisher Scientific, Eindhoven, The Netherlands) set to an acceleration voltage of 80 kV. Final TEM images were manually improved using GIMP v2.10.32 for contrast and brightness adjustment.

#### Host range data

Previously isolated phages, mentioned in the section ‘*Phage assembly, annotation and comparative genomics’*, were included in the host range analysis. All bacterial isolates targeted by a phage are noted in suppl. table 2. The host range assays were done in several rounds (suppl. table 2). For each round a subset of bacterial isolates and phages were tested. All bacterial host were grown ON in LB at 28 °C and all phage lysates were prepared by amplification on their respective host (supplementary fig. 1), prior to each host range assay. 10 µl of phage lysates in dilutions were spotted on a bacterial lawn, using LB agarose (0.4 % agarose) supplemented with 10 mM CaCl_2_·MgCl_2_, with 100 µl of ON host bacteria. Plates were incubated ON at 28 °C before evaluation of the plate.

### Graphics

Graphics were created with R studio (v4.5.1) (R Core Team, 202) using the packages ggtree (v3.16.3) (Yu, 2022), ggtreeExtra (v1.18.0) (Xu et al., 2021), viridis (v0.6.5), ggplot2 (v4.0.0) (Wickham, 2016) and ggimage (v0.3.4) (Yu, 2017).

## RESULTS

A total of 127 samples of diseased potato tubers and plants were received from Danish potato farmers over a period of three years, resulting in 144 isolates, in which 103 met the criteria set for complete genomes and used for downstream analysis. Samples were used to investigate and evaluate the genomic distribution of soft rot pathogens in Danish potato production. Metadata was collected alongside samples and used to investigate the epidemiology of soft rot *Pectobacteriaceae* in potato production across the country. Isolates from the first year of sampling (2021) were used for bacteriophage isolation.

### *P. brasiliense* and *P. atrosepticum* as the dominating species during the years 2021 - 2023

The majority of soft rot isolates were shown to belong to several *Pectobacterium* species; *P. atrosepticum*, *P. brasiliense*, *P. carotovorum*, *P. parmentieri*, *P. Polaris, P. versatile* and *P. punjabense* (Fig. 1A). We did, however, also isolate other genera than *Pectobacterium* such as *Pseudomonas*, *Acinotibacter, Chryseobacterium* and *Serratia*, which were considered false positives (Fig. 1A, species grey). Closely related isolates did in some cases share isolation year and sampling location (Region), sample origin or month. Nevertheless, we observed several closely related strains of some species across sampling locations (Region) as well as both sampling month and year (specified in the next section ‘*Clonal clades of Pectobacterium isolates found across locations and years*’). Across all three years, *P. brasiliense* and *P. atrosepticum* were the most frequently identified isolates, as well as within in the subset of 2021 and 2023. In 2022 *P. parmentieri* was the most frequently isolated species (Fig. 1B). Most of our samples were received during 2021, with Midtjylland and Sydjylland as the most frequent sampling locations (Fig. 1B). July was seen as the predominant month, where more than three times as many samples were received compared to any other month. Likewise, did two thirds of the isolates originate from the stem of plants, symptomatic with blackleg symptoms, and only one third of the samples originated from tubers displaying soft rot symptoms (Fig. 1B), although this may be influenced by sampling instructions. Metadata was collected with samples describing sampling location (Region and Municipality) and potato variety (Fig. 1C). Sample origin as well as date was noted upon receiving the sample and species were assigned to all isolates post-isolation. A total of 22 potato varieties were collected across all three years, where two varieties (Avenue and Saprodi) accounted for almost one third of all samples received (suppl. table 2).

**Figure 1.**
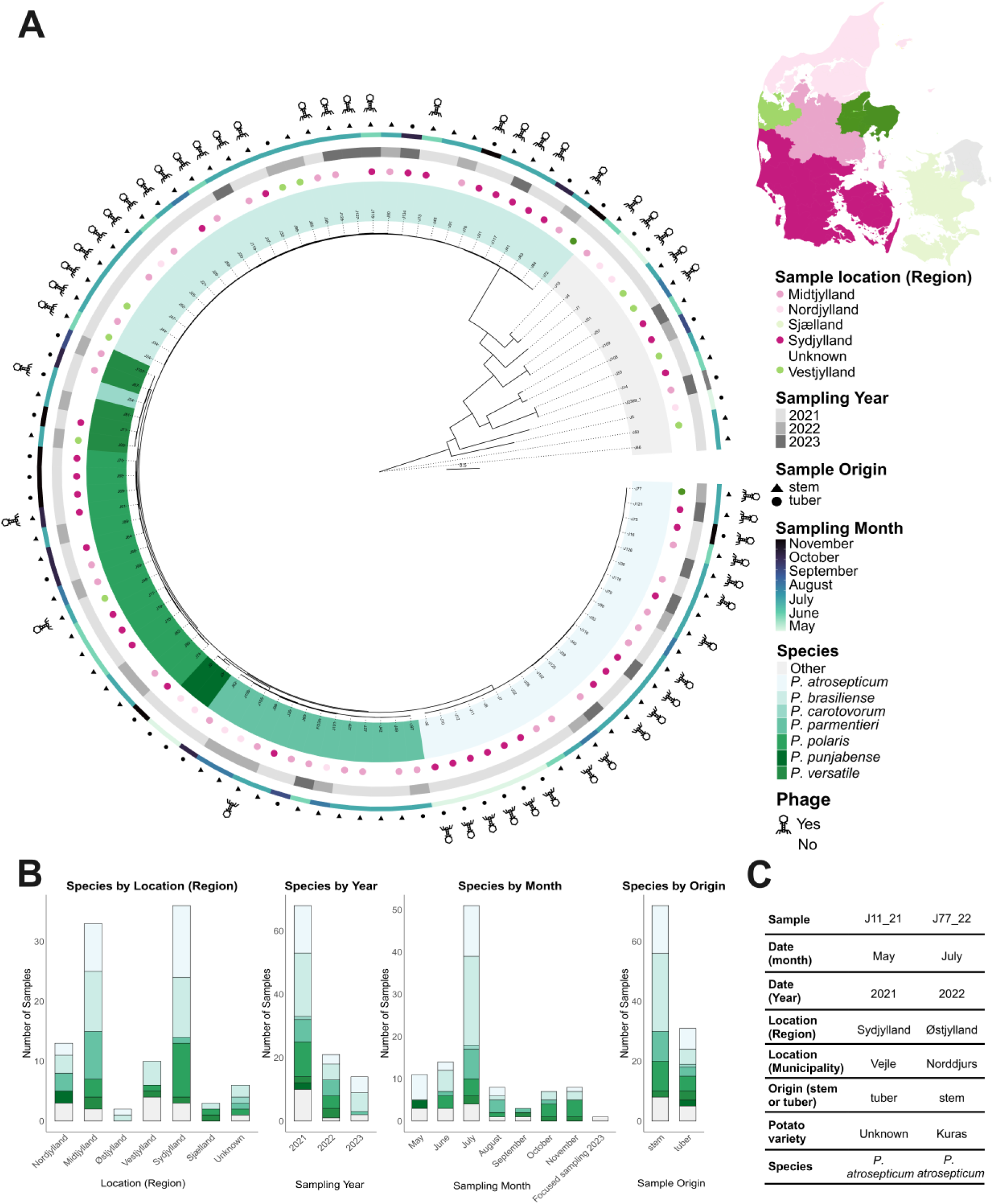
Phylogenetic relationship of all soft rot isolates and corresponding metadata. A) Phylogenetic tree using PhyloPhlAn between 103 Danish soft rot isolates. All isolates are assigned with isolation name and assigned as species, sample location, month, origin and if targeted by a phage. Species by color (1^st^ ring); Other (other than *Pectobacterium* species, light grey) and a colour gradient of green going from light to dark as: *P. atrosepticum*, *P. brasiliense*, *P. carotovoroum*, *P. parmentieri*, *P. Polaris, P. versatile* and *P. punjabense*. Sampling location by colored dots (2^nd^ ring); Midtjylland (rosa), Nordjylland (light rosa), Sjælland (light green), Sydjylland (dark rosa), Unknown (white/no color) and Vestjylland (green). Sampling year by color (3^rd^ ring): 2021 (light grey), 2022 (grey) and 2023 (dark grey). Sampling month by a color gradient, light to dark, as (4^th^ ring): May, June, July, August, September, October and November. Sample origin with square as stem and circle as tuber (5^th^ ring) and phage, implying whether any of the isolated phages are targeting the soft rot isolate, assigned with a phage logo (6^th^ ring). Sampling locations are colored based on region and regions are visualized on Denmark map (upper right corner). B) Species based on sampling location as region, year, month and sample origin. For all plots defined variables within each category are visualized on the x axes (regions, year, month or origin) and a stacked bar plot visualizing the species proportions as number of samples on y axe. All metadata used for the phylogenetic tree are noted in suppl. table 2. C) Example of received metadata and sample details based on two randomly selected samples together with assigned species.

### Clonal clades of *Pectobacterium* isolates found across locations and years

Due to the higher error rate of Nanopore data we only used strains sequenced with Illumina to be included in our SNP analysis, 67 of 103 isolates. Several clonal clades of *Pectobacterium* isolates (defined as less than 20 SNPs detected between strains) were found within *P. atrosepticum*, *P. brasiliense* and *P. polaris* (Fig. 2A). Three clonal clades were found to originate from different samples with <20 SNPs. Two clades of 10 and four, respectively, were found within *P. brasiliense* (*P. brasiliense* clade I and clade II) which for both clades were found across both locations and years. One clade of three, were also found within *P. polaris* (*P. polaris* clade I). *P. Polaris* clade I did originate from two different samples, which shared both location, sampling time and potato variety. Whole genome alignment for all isolates within each clonal clade, to each respective reference, showed genomes within each clade to align across the whole genome (Fig. 2C). Furthermore, did we observe one clonal clade within *P. atrosepticum.* However, *P. atrosepticum* clade I was isolated from the same sample and were assumed to originate from a clonal infection (Suppl. fig. 2).

**Figure 3.**
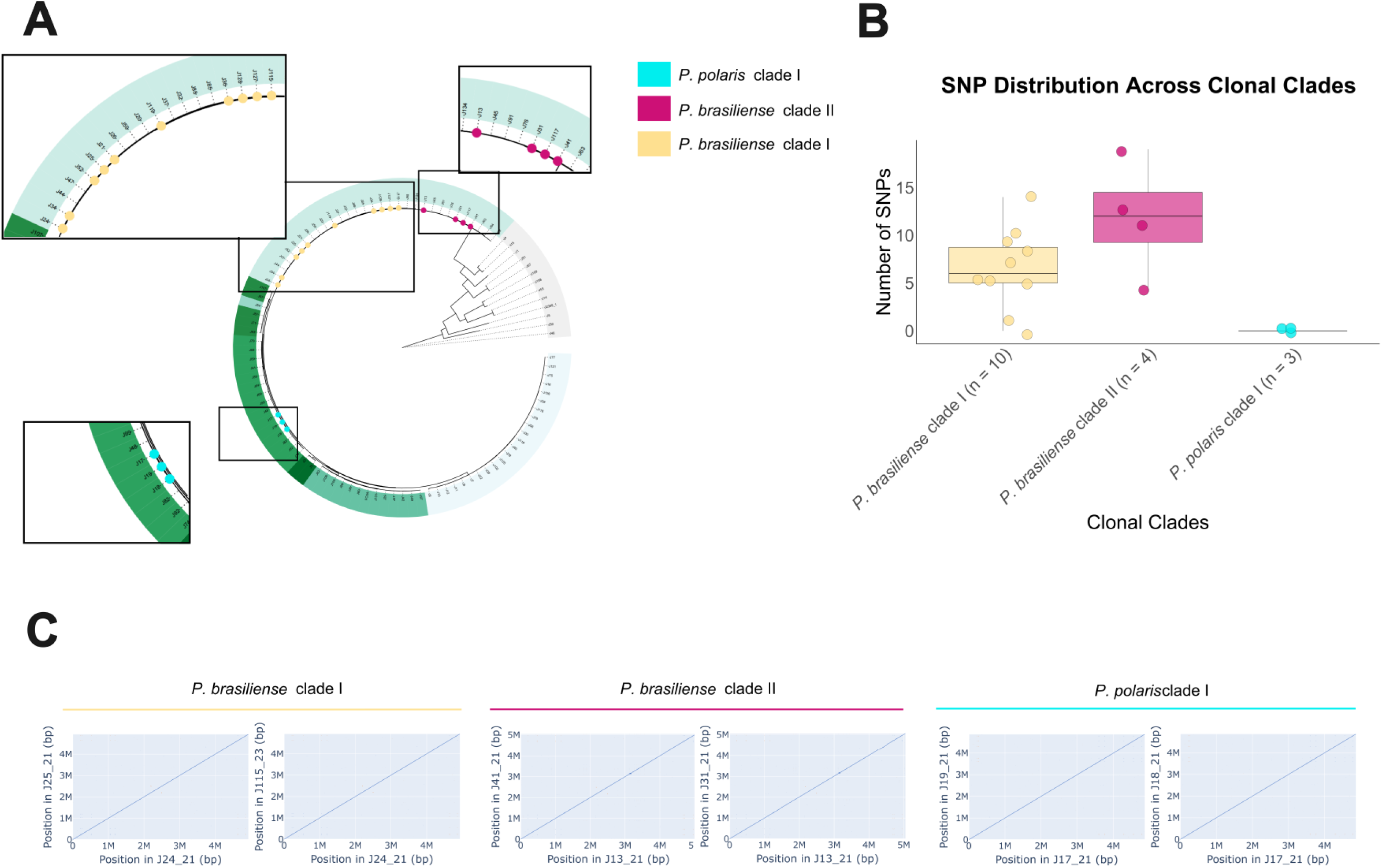
Clonal clades of Pectobacterium isolates. A) Phylogenetic tree assigned with species (as in Fig. 1A), with clonal clades marked as: *P. brasiliense* clade I (yellow), *P. brasiliense* clade II (purple) and *P. polaris* clade I (turquise). B) Single nucleotide polymorphism (SNP) distribution across clonal clades, where number of SNPs are shown on the y axe and clonal clades (including n for each group) on the x axe. Clades are coloured as in A. C) Dotplots visualizing whole genome alignment for two isolates to the reference genome from each clade respectively.

### Isolated phages are targeting most clonal *Pectobacterium* strains

Six phages were isolated using a subset of *Pectobacterium* isolates. These were compared with previously isolated phages targeting *Pectobacterium (Pedersen, Carstens, Bollen, et al., 2025; Pedersen, Carstens, Rothgardt, et al., 2025; Pedersen et al., 2024)* (Fig. 4A, suppl. table 1). The phages showed variation in terms of both genome size and morphology, where most phages had a genome size of 40 to 50 kbp. Notably, phage Magnum possessed the largest genome of ∼ 100 kbp (suppl. table 1). Based on nucleotide similarity two clades were formed, of eight and four phages, respectively. Both clades share >70% nucleotide similarity, representing two genera where phage mspotatoheads were seen to belong to the Ymer genus, previously described (Pedersen et al., 2024). Phage Guf proved part of the second clade, comprised of phage Guf, Crus, Gander and Hoejben (Fig. 4A). The remaining two phages isolated in this study (phage Vims and Magnum) do not group together and share ≤ 2% nucleotide similarity with any of the other phages. Previously isolated phage mrpotatohead and phage Mimer does likewise share ≤ 2% nucleotide similarity with any of the other phages. Transmission electron microscopies were done on three of the six isolated phages (Fig. 4B). Phage Vims showed siphovirus morphology, characterized by long, flexible and non-contractile tails. Phage Vims exhibit multiple small fibers with globular attachments. Phage Guf is a podovirus with a short non-contractile tail. Phage Magnum exhibits a myovirus morphotype with a non-flexible, contractile tail and multiple long tail fibres. We also isolated a total of 9 phages targeting non *Pectobacterium* isolates, targeting both *Serratia* sp., *Morganella* sp. and *Lelliolita* sp. (Suppl. fig. 3, suppl. table 1). The host range results based on phages and soft rot isolates, proved ∼50% of all *Pectobacterium* isolates to be infected by at least one phage, however, most (>80%) of the clonal clades were infected by at least one phage (Fig. 1A (6^th^ ring, phage logo), suppl. table 2). Furthermore, did nine of the phages target at least one or more isolate from the species’ *P. atrosepticum*, *P. brasiliense*, *P. parmentieri* and *P. polaris* (suppl. table 2).

**Figure 4.**
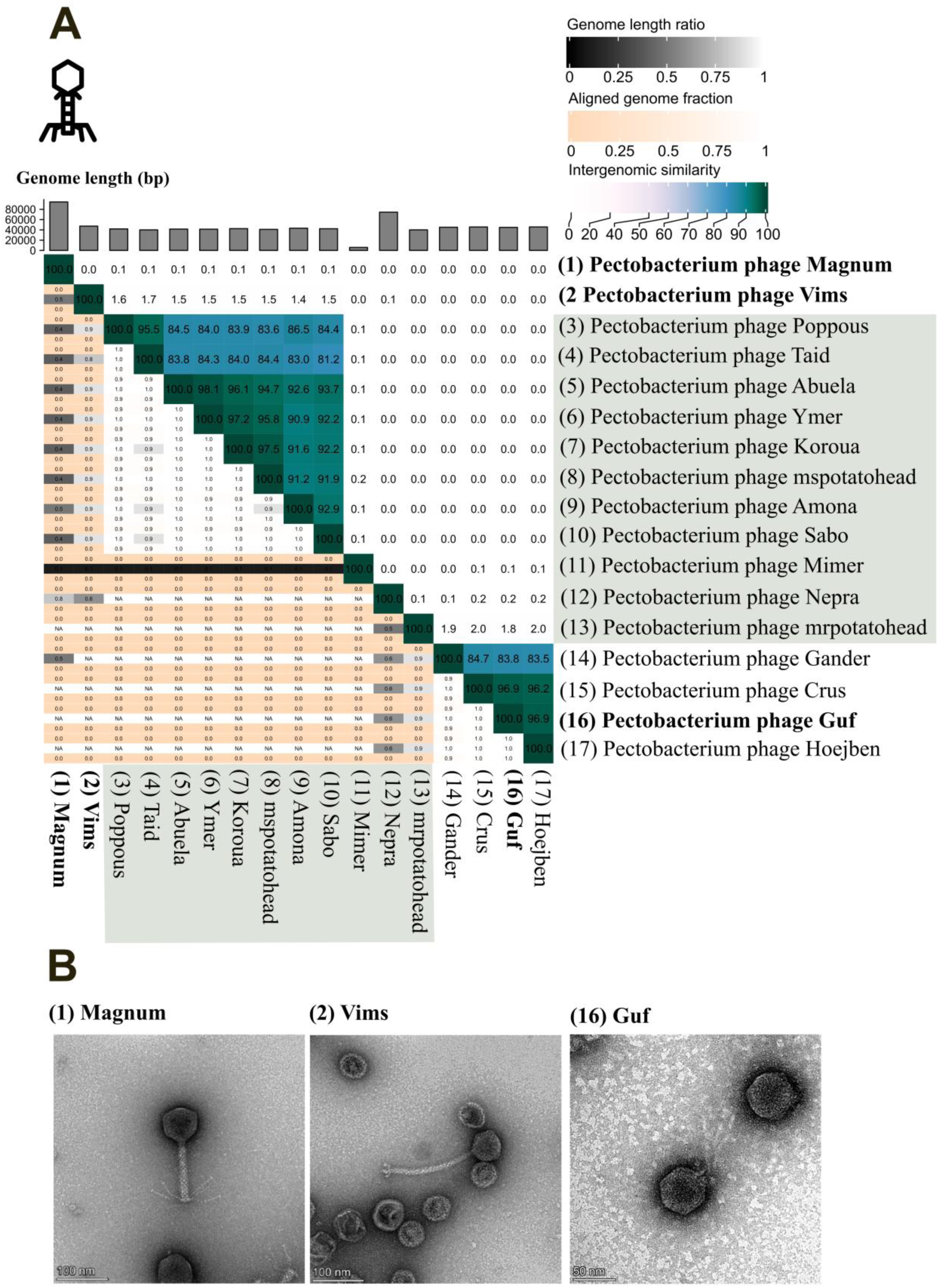
Pectobacterium phages isolates in this study compared with previously isolated phages (A. B. Carstens et al., 2019; Pedersen, Carstens, Bollen, et al., 2025; Pedersen et al., 2024, 2026). A) VIRIDIC heatmap visualizing the intergenomic similarity score of all versus all phages. Genome length (bp) is shown as bars on top of heatmap for each phage, respectively. Aligned genome fraction for each phage in each alignment is shown as a color gradient from from 0 (sand color) to 100 (white). The intergenomic similarity score is shown as a color gradient from low (light) to high (dark) and assigned with a score (0-100) for all alignments. Highlighted phages were previously isolated (suppl. table 1) B) Transmission electron microscopy of three selected Pectobacterium phages. All three phages are tailed and belong to the *Caudoviricetes* class. Phage Vims display a siphovirus morphotype, while phages Guf and Magnum exhibit typical Podovirus and Myovirus morphologies, respectively.

## DISCUSSION

As far as we are aware, this is the first study proving clonal spread of *Pectobacterium* species using WGS and SNP analysis across sample locations and year. Using an agri-citizen science approach, we received tubers and plants symptomatic with soft rot and black leg symptoms from Danish potato farmers over a period of three years. Based on all three years we showed *P. atrosepticum*, *P. brasiliense*, *P. carotovorum*, *P. parmentieri*, *P. Polaris, P. versatile* and *P. punjabense* to be present in Danish potato plants and tubers with disease symptoms, with *P. brasiliense* and *P. atrosepticum* as the most prevalent soft rot species in Denmark. Five different species have been reported in Denmark previously; *P. parmentieri*, *P. atrosepticum*, *P. brasiliense*, *D. solani* and *D. dianthicola* (Ravnskov, 2021). We did isolate *P. parmentieri*, *P. atrosepticum* and *P. brasiliense* across all three years, but did only once isolate *D. dianthicola* and *D. solani*, respectively, (not part of the phylogenetic analysis due to low quality sequencing data, data not shown) across all three years. These results indicate both *Dickeya* species to be less relevant in the Danish potato production compared to prior observations. Interestingly, we also isolated *P. Polaris, P. versatile* and *P. punjabense* which have not been registered in Denmark previously. Ravnskov (2021) did however report the possibility of both *P. polaris* and *P. punjabense* to be present in Denmark, as these had been reported in Norway, the Netherlands and Pakistan.

Three clonal clades (SNP< 20) were found within *Pectobacterium* isolates which varied across samples, locations and/or year (Fig. 2AB). Two clades of *P. brasiliense* included isolates which varied across year and sampling location. *P. brasiliense* clade I did include isolates for two years of sampling, which is a strong indication of a shared infection entry across years. The lag of an apparent relation between sample geographical location and the phylogeny of the isolates combined with the observation of clonal strains isolated at different locations and timepoints, is consistent with contaminated seed or preseed tubers as the source of the inoculum, which could represent an early infection site. Whole-genome sequencing (WGS) is an emerging method to reveal epidemiological patterns in disease transmission, which is increasingly used in agriculture (Weisberg et al., 2021). WGS have been used for studies on the epidemiological spread of SRPs, however mainly focusing on *Dickeya* species which are known to highly clonal (X. Ma et al., 2024; Van Gijsegem et al., 2023). X. Ma et al. studied the epidemiological spread of SRPs across the US of both *Pectobacterium* and *Dickeya* species, using isolates across numerous locations, sites and years. However, only on clade of *D. dianthicola* were proved to be clonal across locations. In relation to clonal spread of other bacterial plant pathogens using WGS and SNP analysis, did Abrahamian et al. suggest bacterial spot on tomatoes, caused by *Xanthomonas perforans* to originate in commercial facilities used for propagation of plants prior to field planting, by genome-wide SNP (Abrahamian et al., 2019). Likewise, our results with the presentation of at least four clonal clades strongly indicate the introduction of *Pectobacterium* at an early stage which could be during the propagation process. One clonal clade was also observed for *P. atrosepticum*, however, *P. atrosepticum* clade I with <20 SNPs were all isolated from the same sample and were assumed to originate from same pathogen (Suppl. fig. 2).

Besides *Pectobacterium* isolates, we also isolated some non-*Pectobacterium* species (Fig 1A, suppl. table 1). All non-*Pectobacterium* species isolates are known to be found in soil, except for *Chishuiella* sp. which has been isolated from fresh water (Ashelford, 2002; Jeon et al., 2024; Mwanza et al., 2022; Ouellette & Wang, 2023; Papade et al., 2024; Sundararaman et al., 2017; Zhang et al., 2014). Nonetheless, few of the non-*Pectobacterium* species, *P. marginalis*, *Lelliottia* sp. and *Chryseobacterium* are described as soil bacteria but also identified as causing soft rot in plants (Liao et al., 1997; Osei et al., 2022; Radke et al., 2024). However, all non-*Pectobacterium* species were only isolated 1-2 times, respectively, during all three years, apart from *Chryseobacterium* which was isolated six times in total. Radke et al. (2024) is the only report on *Chryseobacterium* strains causing soft rot in potato tubers, but *Chryseobacterium* has previously been reported to have a high pectinase activity (Roy et al., 2018). *Chryseobacterium* isolates could be a result of our isolation protocol as this method utilizes pectin degradation as a selective trait. However, we cannot rule out the possibility that these isolates are true soft rot pathogens, as Koch’s postulates were not tested as part of this study.

The mother tuber has previously been suggested as the primary source of infection and considering the post-harvest economic impact of SRPs, post-harvest tuber management represents a critical timepoint for reducing the pathogen load (Dupuis et al., 2021; van der Wolf et al., 2021). Phages have been proposed as a post-harvest control strategy (Thi Trang Nhung et al., 2025). Phages isolated as part of this study together with previously isolated phages did target most (>80%) of the clonal strains, which suggest these to be promising biocontrol agents for application early during the propagation process (seed or pre-seed tubers). Considering all *Pectobacterium* strains isolated across all three years, phages were able to target ∼50% of these. Pectobacterium phages presented in this study together with previously isolated phages may be important in a biocontrol aspect, targeting Danish soft rot *Pectobacterium* strains known to cause disease in the Danish potato production.

## CONCLUSION

Based on the study presented here, we conclude that:

a. Novel and updated knowledge on the prevalence of soft rot species found in Denmark are presented. The genomic dataset can be used for an optimization of SRP screening methods as new primers may be beneficial to uncover the prevalence of these species.
b. Clonal clades and large shifts in species abundance over time (*Dickeya* have seen to disappear and new species have been observed) indicate strong, recent bottlenecks determining SRP prevalence in potato production. It remains unclear what causes these bottlenecks.
c. Clonal clades within more SRP species underscore the need for further research in the pre-seed production to determine if early propagation practices can be improved to limit SRP transmission.
d. As biocontrol management has been pointed towards being more successful if the management window is rather limited, we propose the use of phages as biocontrol agents in the pre-seed production, which may be more impactful than later in the tuber production. However, more research is needed to elucidate these aspects.
e. As phages were able to target most (>80%) of the clonal strains and one third of all *Pectobacterium* strains, further work is needed to establish a cocktail to cover the entire diversity of Danish SRPs.

## ACKNOWLEDGEMENT

This study was funded by the Novo Nordisk Project Grant, grant number: NNF20OC0064283. All authors declare no conflict of interest. We want to show our gratitude to all the Danish potato farmers who supported our project in both 2021, 2022 and 2023. Thanks to KMC Agro, especially Lasse Primdal and Lars Bødker from SEGES for collaboration, idea generation and sharing their huge knowledge on soft rot disease in potatoes and the Danish potato industry during all three years.

## SUPPLEMENTARY MATERIALS

**Supplementary figure 1.**
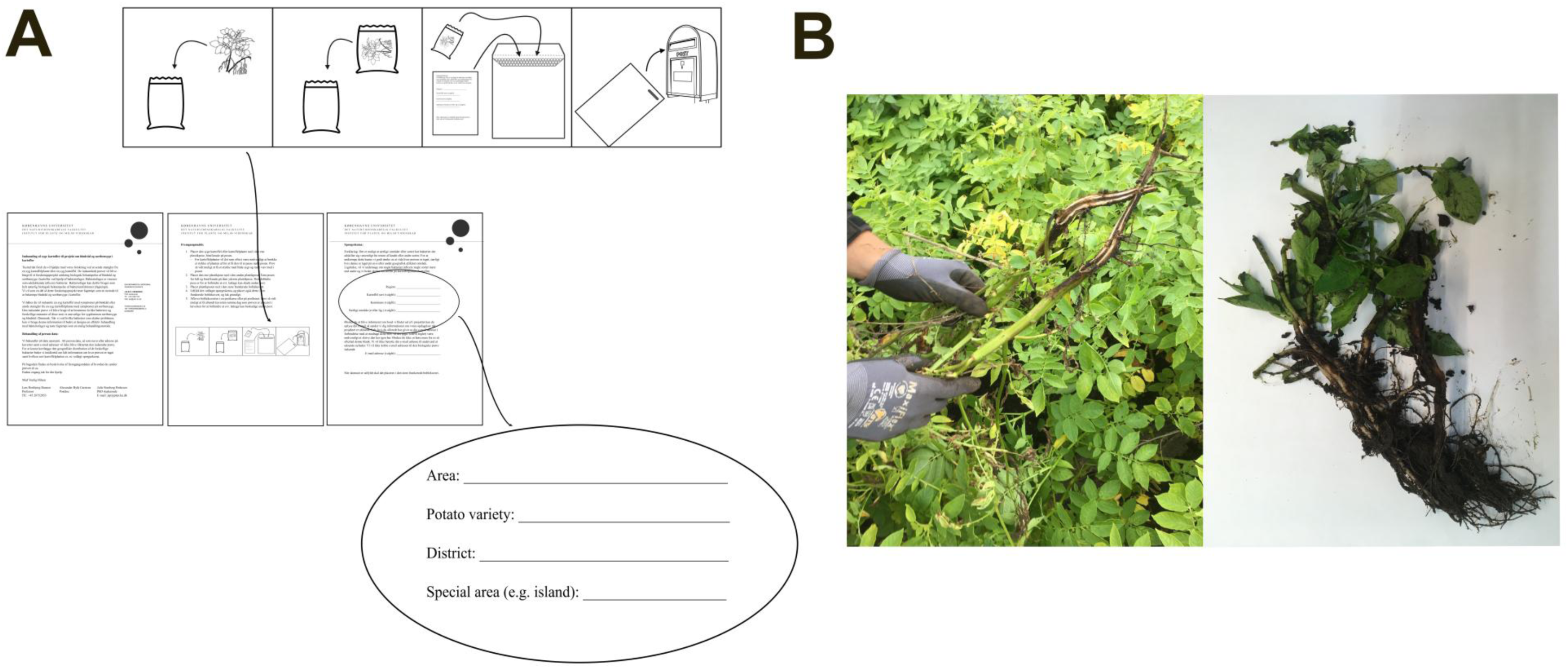
Sample kit for potato farmers across Denmark and soft rot disease symptoms in a plant in a potato field and a received sample. A) Overview of sample kit handed out to potato farmers across Denmark. The sample kit provides a project description and a description on how to use the kit to collect diseased potato plants or tubers and how fill out the accompanying document. In the accompanying document the potato farmer should fill out: area (county), district (municipality), potato variety and if relevant any special area (e.g. island or other). B) Standard soft rot symptoms in a potato plant in the field (left), where part of the stem is turning black. Received sample displaying soft rot symptoms (right), where the stem is black and appear macerated.

**Supplementary figure 2.**
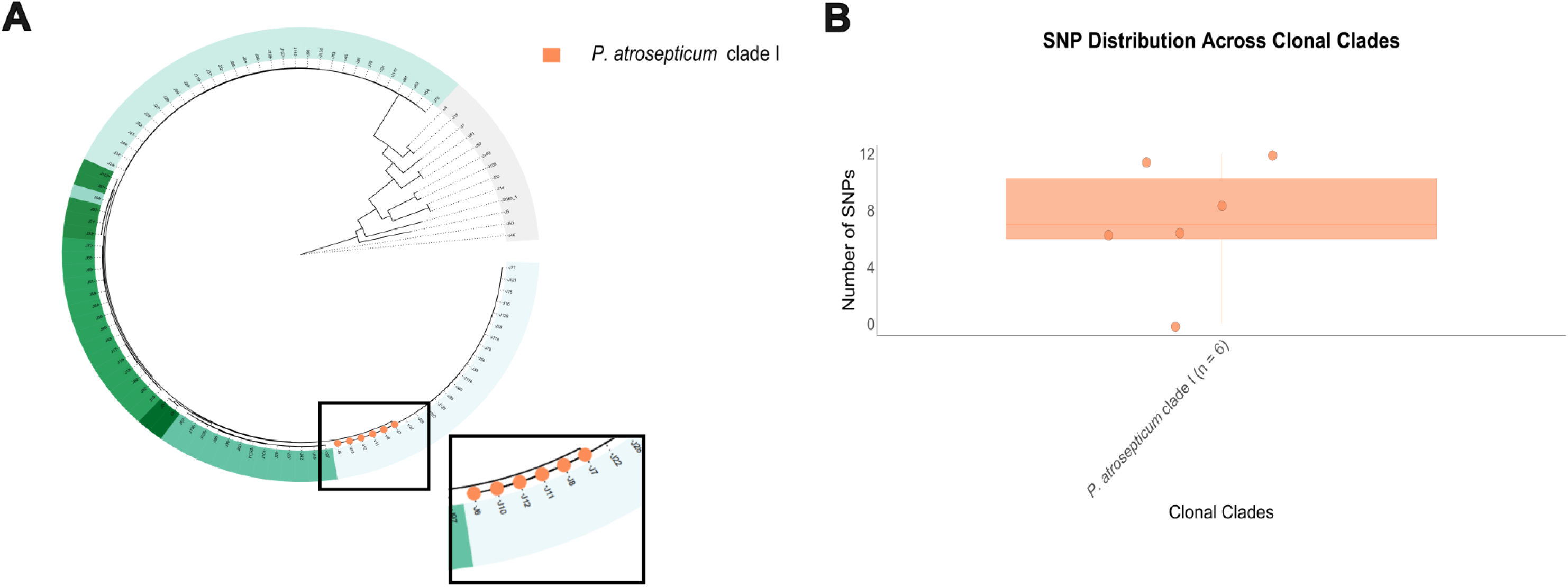
Figure the SNP distribution within one clonal clade of *P. atrosepticum*. A) Phylogenetic tree assigned with species (as in Fig. 1A), with *P. atrosepticum* clade I marked with orange. B) SNP distribution within one clonal clade of *P. atrosepticum*. Clade I includes 6 isolates with (SNPs < 20), all isolated from the same sample.

**Supplementary table 1.** Overview of all isolated phages. Pectobacterium phages are numbered as seen in fig. 4A and non-Pectobacterium phages are numbered as in Suppl. fig. 1. Phages are assigned with GenBank ass. no. (if available), Reference (or this study), whether it is a new genus or not, genome size, host ID, host species and target species in the host range analysis.

| Phage | GenBank acc. no. | Reference | New genus (yes/no) | Genome size (bp) | Host ID | Host Species | Target species |
| --- | --- | --- | --- | --- | --- | --- | --- |
| (1) Pectobacterium phage Magnum | - | this study | no | 94385 | J48_21 | *P. polaris* | *P. polaris*, *P. atrosepticum*, *P. brasiliense*, *P. parmentieri* |
| (2) Pectobacterium phage Vims | - | this study | yes | 47310 | J58_21 | *P. parmentieri* | *P. parmentieri* |
| (3) Pectobacterium phage Poppous | PQ008973 | Pedersen et al., 2024 | (Ymer genus) | 40850 | J13_21 | *P. brasiliense* | *P. brasiliense*, *P. parmentieri*, *P. polaris* |
| (4) Pectobacterium phage Taid | PQ008975 | Pedersen et al., 2024 | (Ymer genus) | 39941 | J31_21 | *P. brasiliense* | *P. brasiliense*, *P. parmentieri*, *P. polaris*, *P. atrosepticum* |
| (5) Pectobacterium phage Abuela | OP748251 | Pedersen et al., 2024 | (Ymer genus) | 41492 | J34_21 | *P. brasiliense* | *P. brasiliense*, *P. parmentieri*, *P. polaris* |
| (6) Pectobacterium phage Ymer | OP748252 | Pedersen et al., 2024 | (Ymer genus) | 41279 | J36_21 | *P. brasiliense* | *P. brasiliense*, *P. parmentieri*, *P. polaris*, *P. atrosepticum* |
| (7) Pectobacterium phage Koroua | PQ008972 | Pedersen et al., 2024 | (Ymer genus) | 41346 | J52_21 | *P. brasiliense* | *P. brasiliense*, *P. parmentieri* |
| (8) Pectobacterium phage mspotatohead | - | this study | no | 40933 | J47_21 | *P. brasiliense* | not tested |
| (9) Pectobacterium phage Amona | PQ008971 | Pedersen et al., 2024 | (Ymer genus) | 42206 | J25_21 | *P. brasiliense* | *P. brasiliense*, *P. parmentieri*, *P. polaris*, *P. atrosepticum* |
| (10) Pectobacterium phage Sabo | PQ008974 | Pedersen et al., 2024 | (Ymer genus) | 42167 | J24_21 | *P. brasiliense* | *P. brasiliense* |
| (11) Pectobacterium phage Mimer | ON872163 | Pedersen et al., 2026 | yes | 5802 | J47_21 | *P. brasiliense* | *P. brasiliense*, *P. parmentieri*, *P. polaris*, *P. atrosepticum* |
| (12) Pectobacterium Nepra | NC_048704.1 | A. B. Carstens et al., 2019 | no | 74509 | DSM18077 | *P. atrosepticum* | *P. atrosepticum* |
| (13) Pectobacterium phage mrpotatohead | - | this study | no | 40065 | J47_21 | *P. brasiliense* | not tested |
| (14) Pectobacterium phage Gander | - | this study | no | 45139 | J44_21 | *P. brasiliense* | *P. brasiliense*, *P. parmentieri*, *P. polaris*, *P. atrosepticum* |
| (15) Pectobacterium phage Crus | - | this study | no | 45644 | J36_21 | *P. brasiliense* | *P. brasiliense*, *P. parmentieri*, *P. polaris*, *P. atrosepticum* |
| (16) Pectobacterium phage Guf | - | this study | no | 44908 | J26_21 | *P. brasiliense* | *P. brasiliense*, *P. parmentieri*, *P. polaris*, *P. atrosepticum* |
| (17) Pectobacterium phage Hoejben | - | this study | no | 45692 | J32_21 | *P. brasiliense* | *P. brasiliense*, *P. parmentieri*, *P. polaris*, *P. atrosepticum* |
| (1) Serratia phage Uther | - | this study | no | 44796 | J4_21 | *S. plymuthica* | not tested |
| (2) Serratia phage Echoes | - | this study | no | 59690 | J15_21 | *S. liquefaciens* | not tested |
| (3) Morganella phage Rip | - | this study | no | 59319 | J51_21 | *M. morganii* | not tested |
| (4) Morganella phage Rup | - | this study | no | 60952 | J57_21 | *M. morganii* | not tested |
| (5) Lelliottia phage Pantea | - | this study | no | 150009 | J1_21 | *L. amnigena* | not tested |
| (6) Serratia phage Galvinrad | - | this study | no | 32655 | J4_21 | *S. plymuthica* | not tested |
| (7) Morganella phage Slaad | - | this study | no | 43547 | J57_21 | *M. morganii* | not tested |
| (8) Lelliottia phage Rap | - | this study | yes | 40557 | J1_21 | *L. amnigena* | *L. amnigena* |
| (9) Lelliottia phage Zann | - | this study | no | 51744 | J1_22 | *L. amnigena* | *L. amnigena* |

**Supplementary table 2.** Overview of all bacterial isolates used for phylogenetic analysis together with assigned metadata. All isolates are assigned with species, sample location (Region), sample location (Municipality), sampling month, sampling year, sample origin, potato variety and phage (whether any phages were able to infect the isolate in the host range analysis.

| ID | Species | Location (Region) | Location (Municipality) | Sampling month | Sampling year | Sample ori. | Potato var. | Phage |
| --- | --- | --- | --- | --- | --- | --- | --- | --- |
| FO3A | *Pectobacterium parmentieri* | Midtjylland | Unknown | June | 2023 | stem | Saprodi |  |
| J101 | *Pectobacterium parmentieri* | Nordjylland | Frederikshavn | August | 2022 | stem | Saprodi |  |
| J102 | *Pectobacterium atrosepticum* | Nordjylland | Jammerbugt | August | 2022 | stem | Ydun | + |
| J10 | *Pectobacterium atrosepticum* | Syddjylland | Vejle | May | 2021 | tuber | Unknown | + |
| J105 | *Pectobacterium parmentieri* | Nordjylland | Hjørring | August | 2022 | stem | Ydun |  |
| J106 | *Pectobacterium parmentieri* | Nordjylland | Hjørring | August | 2022 | stem | Ydun |  |
| J107 | *Pectobacterium versatile* | Midtjylland | Skive | September | 2022 | tuber | Avenue |  |
| J108 | *Pseudomonas marginalis* | Syddjylland | Varde | September | 2022 | tuber | Colombo |  |
| J109 | *Pseudomonas chlororaphis* | Syddjylland | Varde | June | 2023 | stem | Ydun |  |
| J11 | *Pectobacterium atrosepticum* | Syddjylland | Vejle | May | 2021 | tuber | Unknown | + |
| J115 | *Pectobacterium brasiliense* | Syddjylland | Ringkøbing-Skjern | June | 2023 | stem | Saprodi | + |
| J116 | *Pectobacterium atrosepticum* | Syddjylland | Ringkøbing-skjern | July | 2023 | stem | Ydun | + |
| J118 | *Pectobacterium atrosepticum* | Syddjylland | Billund | July | 2023 | stem | Fyone |  |
| J117 | *Pectobacterium brasiliense* | Syddjylland | Haderslev | July | 2023 | stem | Saturna | + |
| J119 | *Pectobacterium brasiliense* | Nordjylland | Thisted | July | 2023 | stem | Spunta | + |
| J121 | *Pectobacterium atrosepticum* | Syddjylland | Ringkøbing-Skjern | July | 2023 | stem | Avenue | + |
| J125 | *Pectobacterium atrosepticum* | Midtjylland | Ikast-Brande | July | 2023 | stem | Stratos |  |
| J126 | *Pectobacterium atrosepticum* | Nordjylland | Frederikshavn | July | 2023 | stem | Ydun | + |
| J127 | *Pectobacterium brasiliense* | Nordjylland | Frederikshavn | July | 2023 | stem | Avenue | + |
| J128 | *Pectobacterium brasiliense* | Midtjylland | Skive | July | 2023 | stem | Verdi | + |
| J12 | *Pectobacterium atrosepticum* | Syddjylland | Vejle | May | 2021 | tuber | Unknown | + |
| J134 | *Pectobacterium brasiliense* | Syddjylland | Vejen | October | 2023 | tuber | Avenue |  |
| J13 | *Pectobacterium brasiliense* | Syddjylland | Billund | June | 2021 | stem | Kardal | + |
| J14 | *Acinetobacter calcoaceticus* | Syddjylland | Billund | June | 2021 | stem | Kardal |  |
| J15 | *Serratia liquefaciens* | Midtjylland | Ikast-Brande | June | 2021 | stem | Kuras | + |
| J16 | *Pectobacterium atrosepticum* | Midtjylland | Ikast-Brande | June | 2021 | stem | Kuras | + |
| J17 | *Pectobacterium polaris* | Syddjylland | Billund | June | 2021 | stem | Kardal |  |
| J18 | *Pectobacterium polaris* | Syddjylland | Billund | June | 2021 | stem | Kardal |  |
| J19 | *Pectobacterium polaris* | Syddjylland | Billund | June | 2021 | stem | Kardal |  |
| J1 | *Lelliottia* | Nordjylland | Vesterhimmerlands | May | 2021 | tuber | Twister | + |
| J20 | *Pectobacterium brasiliense* | Midtjylland | Brande | June | 2021 | stem | Saprodi | + |
| J21 | *Pectobacterium brasiliense* | Midtjylland | Brande | June | 2021 | stem | Saprodi | + |
| J22 | *Pectobacterium atrosepticum* | Midtjylland | Silkeborg | June | 2021 | stem | Kuras |  |
| J24 | *Pectobacterium brasiliense* | Vestjylland | Unknown | June | 2021 | stem | Solist | + |
| J25 | *Pectobacterium brasiliense* | Nordjylland | Aalborg | July | 2021 | stem | Wutan | + |
| J26 | *Pectobacterium brasiliense* | Sjælland | Lolland | July | 2021 | stem | Tyson | + |
| J27 | *Pectobacterium parmentieri* | Midtjylland | Ikast-Brande | July | 2021 | stem | Kuras |  |
| J28 | *Pectobacterium atrosepticum* | Midtjylland | Skjern | July | 2021 | stem | Avenue | + |
| J29 | *Pectobacterium parmentieri* | Midtjylland | Skjern | July | 2021 | stem | Kuras |  |
| J2 | *Pectobacterium punjabense* | Nordjylland | Vesterhimmerlands | May | 2021 | tuber | Twister |  |
| J30 | *Pectobacterium parmentieri* | Syddjylland | Vejle | July | 2021 | stem | Kardal |  |
| J31 | *Pectobacterium brasiliense* | Syddjylland | Billund | July | 2021 | stem | Avenue | + |
| J32 | *Pectobacterium brasiliense* | Syddjylland | Ringkøbing-Skjern | July | 2021 | stem | Frig | + |
| J33 | *Pectobacterium atrosepticum* | Midtjylland | Herning | July | 2021 | stem | Kuras | + |
| J34 | *Pectobacterium brasiliense* | Midtjylland | Skive | July | 2021 | stem | Kardal | + |
| J36 | *Pectobacterium brasiliense* | Midtjylland | Herning | July | 2021 | stem | Ydun | + |
| J37 | *Pectobacterium brasiliense* | Midtjylland | Herning | July | 2021 | stem | Avarna | + |
| J38 | *Pectobacterium atrosepticum* | Midtjylland | Ikast-Brande | July | 2021 | stem | Avarna | + |
| J39 | *Pectobacterium atrosepticum* | Syddjylland | Vejle | July | 2021 | stem | Saprodi | + |
| J3 | *Pectobacterium punjabense* | Nordjylland | Vesterhimmerlands | May | 2021 | tuber | Twister |  |
| J40 | *Pectobacterium atrosepticum* | Syddjylland | Vejle | July | 2021 | stem | Stratos | + |
| J41 | *Pectobacterium brasiliense* | Syddjylland | Ringkøbing-Skjern | July | 2021 | stem | Cimega | + |
| J42 | *Pectobacterium parmentieri* | Unknown | Unknown | July | 2021 | stem | Mikado |  |
| J44 | *Pectobacterium brasiliense* | Vestjylland | Holstebro | July | 2021 | stem | Avenue | + |
| J45 | *Pectobacterium brasiliense* | Unknown | Unknown | July | 2021 | stem | Mikado |  |
| J46 | *Chryseobacterium* | Unknown | Unknown | July | 2021 | stem | Verdi |  |
| J47 | *Pectobacterium brasiliense* | Unknown | Unknown | July | 2021 | stem | Verdi | + |
| J48 | *Pectobacterium polaris* | Vestjylland | Holstebro | July | 2021 | stem | Kardal | + |
| J49 | *Pectobacterium parmentieri* | Midtjylland | Viborg | July | 2021 | stem | Kardal |  |
| J4 | *Serratia plymuthica* | Nordjylland | Vesterhimmerlands | May | 2021 | tuber | Twister | + |
| J50 | *Chryseobacterium* | Vestjylland | Holstebro | July | 2021 | stem | Verdi |  |
| J51 | *Morganella morganii* | Vestjylland | Holstebro | July | 2021 | stem | Avenue | + |
| J52 | *Pectobacterium brasiliense* | Midtjylland | Viborg | July | 2021 | stem | Avenance | + |
| J53 | *Acinetobacter calcoaceticus* | Vestjylland | Holstebro | July | 2021 | stem | Arielle |  |
| J54 | *Pectobacterium carotovorum* | Unknown | Unknown | July | 2021 | tuber | Solist | + |
| J56 | *Pectobacterium atrosepticum* | Midtjylland | Skjern | August | 2021 | stem | Avenue |  |
| J57 | *Morganella morganii* | Vestjylland | Holstebro | August | 2021 | stem | Avenue | + |
| J59 | *Pectobacterium brasiliense* | Syddjylland | Vejen | August | 2021 | stem | Saprodi | + |
| J5 | *Vagococcus* | Nordjylland | Vesterhimmerlands | May | 2021 | tuber | Twister |  |
| J60 | *Pectobacterium parmentieri* | Midtjylland | Skive | September | 2021 | tuber | Avenue |  |
| J6 | *Pectobacterium atrosepticum* | Syddjylland | Vejle | May | 2021 | tuber | Unknown | + |
| J62 | *Pectobacterium parmentieri* | Midtjylland | Herning | October | 2021 | tuber | Kuras |  |
| J63 | *Pectobacterium brasiliense* | Syddjylland | Tønder | October | 2021 | tuber | Avenue |  |
| J64 | *Pectobacterium polaris* | Syddjylland | Haderslev | October | 2021 | stem | Kuras |  |
| J61 | *Pectobacterium polaris* | Syddjylland | Vejen | October | 2021 | stem | Avenue | + |
| J66 | *Pectobacterium polaris* | Midtjylland | Herning | October | 2021 | tuber | Avenue |  |
| J67 | *Pectobacterium versatile* | Midtjylland | Herning | October | 2021 | tuber | Verdi |  |
| J68 | *Pectobacterium polaris* | Syddjylland | Vejle | November | 2021 | tuber | Unknown |  |
| J69 | *Pectobacterium polaris* | Syddjylland | Vejle | November | 2021 | tuber | Unknown |  |
| J70 | *Pectobacterium polaris* | Syddjylland | Vejle | November | 2021 | tuber | Unknown |  |
| J71 | *Pectobacterium versatile* | Syddjylland | Vejle | November | 2021 | tuber | Unknown |  |
| J72 | *Pectobacterium brasiliense* | Østjylland | Norddjurs | November | 2021 | tuber | Saprodi |  |
| J74 | *Pectobacterium polaris* | Syddjylland | Vejen | November | 2021 | tuber | Saprodi |  |
| J7 | *Pectobacterium atrosepticum* | Syddjylland | Vejele | May | 2021 | tuber | Unknown | + |
| J75 | *Pectobacterium atrosepticum* | Syddjylland | Haderslev | November | 2021 | tuber | Avenue | + |
| J76 | *Pectobacterium brasiliense* | Syddjylland | Ringkøbing-Skjern | November | 2021 | tuber | Unknown | + |
| J77 | *Pectobacterium atrosepticum* | Østjylland | Norrdjurs | July | 2022 | stem | Kuras | + |
| J79 | *Pectobacterium atrosepticum* | Midtjylland | Viborg | July | 2022 | stem | Kuras | + |
| J81 | *Pectobacterium versatile* | Sjælland | Lolland | July | 2022 | stem | Tyson |  |
| J82 | *Pectobacterium polaris* | Sjælland | Lolland | July | 2022 | stem | Spunta |  |
| J8 | *Pectobacterium atrosepticum* | Syddjylland | Vejele | May | 2021 | tuber | Unknown | + |
| J84 | *Pectobacterium brasiliense* | Midtjylland | Viborg | July | 2022 | stem | Solist | + |
| J85 | *Pectobacterium brasiliense* | Vestjylland | Holstebro | July | 2022 | stem | Kurdal |  |
| J88 | *Pectobacterium brasiliense* | Vestjylland | Holstebro | July | 2022 | tuber | Ydun |  |
| J89 | *Pectobacterium polaris* | Unknown | Unknown | July | 2022 | stem |  |  |
| J90 | *Pectobacterium brasiliense* | Midtjylland | Holstebro | July | 2022 | stem | Tarzan |  |
| J91 | *Pectobacterium brasiliense* | Midtjylland | Skive | July | 2022 | stem | Verdi |  |
| J92 | *Pectobacterium polaris* | Midtjylland | Ikast-Brande | July | 2022 | stem | Kardal |  |
| J93 | *Pectobacterium versatile* | Vestjylland | Holstebro | July | 2022 | stem | Verdi |  |
| J97 | *Pectobacterium parmentieri* | Midtjylland | Herning | July | 2022 | tuber | Saprodi |  |
| J98 | *Pectobacterium parmentieri* | Midtjylland | Viborg | July | 2022 | stem | Avenue | + |
| J99 | *Pectobacterium polaris* | Midtjylland | Herning | August | 2022 | stem | Kuras |  |
| J2369 1 | *Paenarthrobacter ilicis* | Midtjylland | Unknown | Unknown | 2023 | tuber | Saprodi |  |

**Supplementary figure 3.**
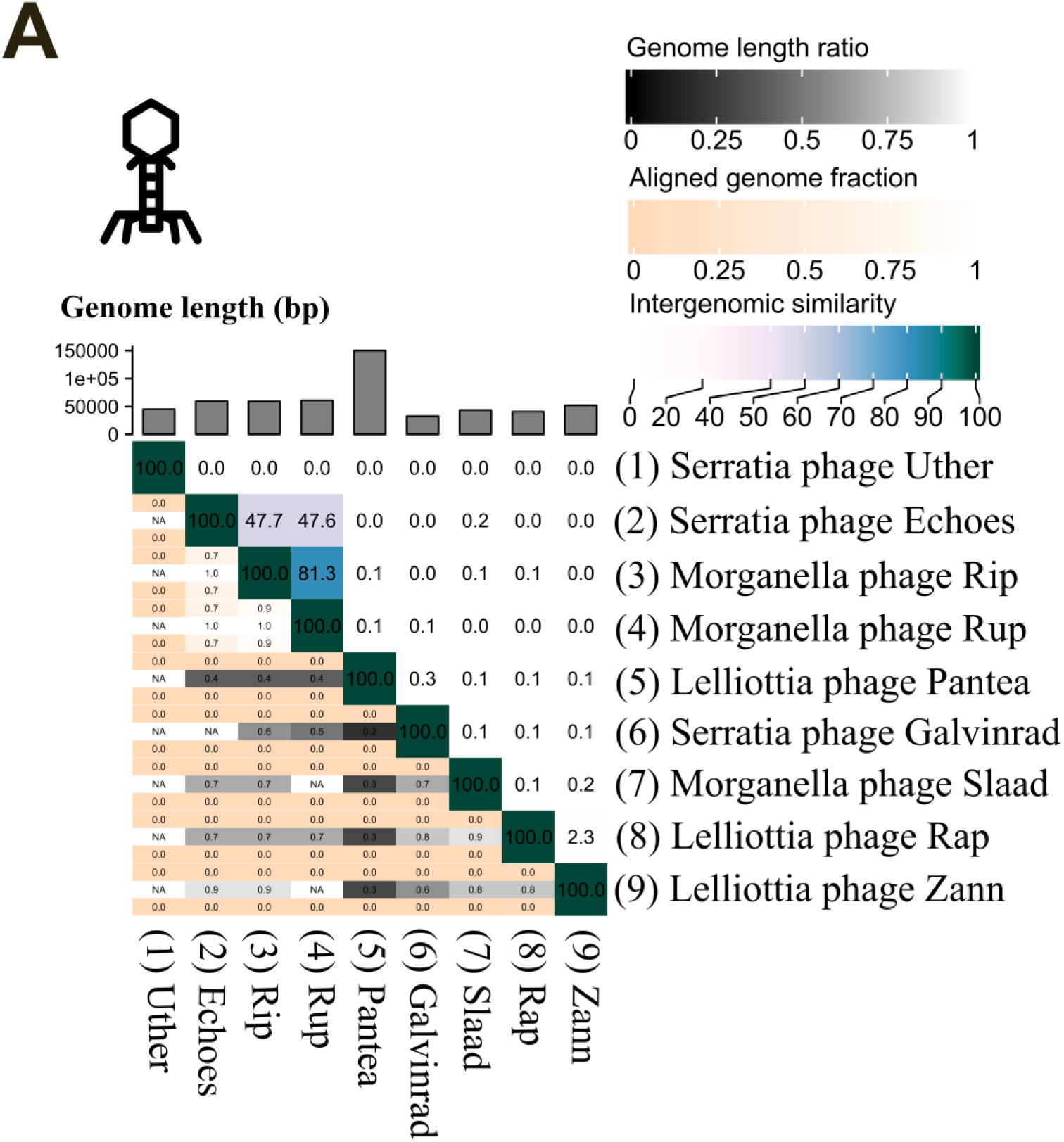
Phages isolated against non-Pectobacterium isolates. VIRIDIC heatmap visualizing the intergenomic similarity score of all versus all phages. Genome length (bp) is shown as bars on top of heatmap for each phage, respectively. Aligned genome fraction for each phage in each alignment is shown as a color gradient from from 0 (sand color) to 100 (white). The intergenomic similarity score is shown as a color gradient from low (light) to high (dark) for all alignments.

